# Alternative splicing links histone modifications to stem cell fate decision

**DOI:** 10.1101/181875

**Authors:** Yungang Xu, Weiling Zhao, Scott D. Olson, Karthik S. Prabhakara, Xiaobo Zhou

## Abstract

**Background:** Understanding the embryonic stem cell (ESC) fate decision between self-renewal and proper differentiation is important for developmental biology and regenerative medicine. Attention has focused on mechanisms involving histone modifications, alternative pre-mRNA splicing, and cell-cycle progression. However, their intricate interrelations and joint contributions to ESC fate decision remain unclear.

**Results:** We analyze the transcriptomes and epigenomes of human ESC and five types of differentiated cells. We identify thousands of alternatively spliced exons and reveal their development and lineage-dependent characterizations. Several histone modifications show dynamic changes in alternatively spliced exons and three are strongly associated with 52.8% of alternative splicing events upon hESC differentiation. The histone modification-associated alternatively spliced genes predominantly function in G2/M phases and ATM/ATR-mediated DNA damage response pathway for cell differentiation, whereas other alternatively spliced genes are enriched in the G1 phase and pathways for self-renewal. These results imply a potential epigenetic mechanism by which some histone modifications contribute to ESC fate decision through the regulation of alternative splicing in specific pathways and cell-cycle genes. Supported by experimental validations and extended dataset from Roadmap/ENCODE projects, we exemplify this mechanism by a cell cycle-related transcription factor, PBX1, which regulates the pluripotency regulatory network by binding to NANOG. We suggest that the isoform switch from PBX1a to PBX1b links H3K36me3 to hESC fate determination through the PSIP1/SRSF1 adaptor, which results in the exon skipping of PBX1.

**Conclusion:** We reveal the mechanism by which alternative splicing links histone modifications to stem cell fate decision.

## Background

Embryonic stem cells (ESCs), the pluripotent stem cells derived from the inner cell mass of a blastocyst, provide a vital tool for studying the regulation of early embryonic development and cell-fate decision, and hold the promise for regenerative medicine [1]. The past few years have witnessed remarkable progress in understanding the ESC fate decision, i.e. either pluripotency maintenance (self-renewal) or proper differentiation [2]. The underlying mechanisms have been largely expanded from the core pluripotent transcription factors (TFs) [3], signaling pathways [4-10], specific microRNAs [11, 12], and long non-coding RNAs [13] to alternative pre-mRNA splicing (AS) [14, 15], histone modifications (HMs) [16-20], and cell-cycle machinery [21]. These emerging mechanisms suggest their intricate interrelations and potential joint contributions to ESC pluripotency and differentiation, which, however, remain unknown.

Alternative splicing (AS) is one of the most important pre-mRNA processing to increase the diversity of transcriptome and proteome in tissue- and development-dependent manners [22]. The estimates based on RNA-seq revealed that up to 94%, 60% and 25% of genes in human, *Drosophila melanogaster* and *Caenorhabditis elegans*, respectively, undergo AS [22-26]. AS also provides a powerful mechanism to control the developmental decision in ESCs [27-29]. Specific isoforms are necessary to maintain both the identity and activity of stem cells and switching to different isoforms ensures proper differentiation [30]. Especially, the AS of TFs plays major roles in ESC fate determination, such as *FGF4* [31] and *FOXP1* [14] for hESC, *Tcf3* [15] and *Sall4* [32] for mouse ESCs (mESCs). Understanding the precise regulations on AS would contribute to the elucidation of ESC fate decision and has attracted extensive efforts [33]. For many years, studies aiming to shed light on this process focused on the RNA level, characterizing the manner by which splicing factors (SFs) and auxiliary proteins interact with splicing signals, thereby enabling, facilitating and regulating RNA splicing. These *cis*-acting RNA elements and trans-acting splicing factors have been assembled into splicing code [34], revealing a number of AS regulators critical for ESC differentiation, such as MBNL [35] and SON [29]. However, these genetic controls are far from sufficient to explain the faithful regulation of AS [36]. Especially in some cases that tissue-specific AS patterns exist despite the identity in sequences and ubiquitously expression of involved SFs [37, 38], indicating additional regulatory layers leading to specific AS patterns. As expected, we are increasingly aware that splicing is not an isolated process; rather it occurs co-transcriptionally and is presumably also regulated by transcription-related processes. Emerging provocative studies have unveiled that AS is subject to extensive controls not only from genetic but also epigenetic mechanisms due to its co-transcriptional occurrence [39]. The epigenetic mechanisms, such as HMs, benefits ESCs by providing an epigenetic memory for splicing decisions so that the splicing pattern could be passed on during self-renewal and be modified during differentiation without the requirement of establishing new AS rules [39].

HMs have long been thought to play crucial roles in ESC maintenance and differentiation by determining what parts of the genome are expressed. Specific genomic regulatory regions, such as enhancers and promoters, undergo dynamic changes in HMs during ESC differentiation to transcriptionally permit or repress the expression of genes required for cell fate decision [40]. For example, the co-occurrence of the active (H3K4me3) and repressive (H3K27me3) HMs at the promoters of developmentally regulated genes defines the bivalent domains, resulting in the poised states of these genes [41]. These poised states will be dissolved upon differentiation to allow these genes to be active or more stably repressed depending on the lineage being specified, which enables the ESCs to change their identities [42]. In addition to above roles in determining transcripts abundance, HMs are emerging as major regulators to define the transcripts structure by determining how the genome is spliced when being transcribed, adding another layer of regulatory complexity beyond the genetic splicing code [43]. A number of HMs, such as H3K4me3 [44], H3K9me3 [45], H3K36me3 [46, 47], and hyperacetylation of H3 and H4 [48-52], have been proven to regulate AS by either directly recruiting chromatin-associated factors and SFs or indirectly modulating transcriptional elongation rate [39]. Together, these studies reveal that HMs determine not only what parts of the genome are expressed, but also how they are spliced. However, few studies focused on the detailed mechanisms, i.e., epigenetic regulations on AS in the context of cell fate decision.

Additionally, cell-cycle machinery dominates the mechanisms underlying ESC pluripotency and differentiation [21, 53]. Changes of cell fates require going through the cell-cycle progression. Studies in mESCs [54] and hESCs [55, 56] found that the cell fate specification starts in G1 phase when ESCs can sense differentiation signals. Cell fate commitment is only achieved in G2/M phases when pluripotency is dissolved through cell-cycle dependent mechanisms. However, whether the HMs and AS and their interrelations are involved in these cell cycle-dependent mechanisms remains unclear. Therefore, it is intuitive to expect that HMs could contribute to ESC pluripotency and differentiation by regulating the AS of genes required for specific processes, such cell-cycle progression. Nevertheless, we do not even have a comprehensive view of how HMs relate to AS outcome at genome-wide during ESC differentiation. Therefore, further studies are required to elucidate the extent to which the HMs are associated with specific splicing repertoire and their joint contributions to ESC fate decision between self-renewal and proper differentiation.

To address these gaps in current knowledge, we performed genome-wide association studies between transcriptome and epigenome of the differentiation from the hESCs (H1 cell line) to five differentiated cell types [16]. These cells cover three germ layers for embryogenesis, adult stem cells, and adult somatic cells, representing multiple lineages of different developmental levels (Additional file 1: Figure S1A). This carefully selected dataset enabled our understanding of AS epigenetic regulations in the context of cell fate decision. First, we identified several thousands of AS events that are differentially spliced between the hESCs and differentiated cells, including 3,513 mutually exclusive exons (MXE) and 3,678 skipped exons (SE) which were used for further analyses. These hESC differentiation-related AS events involve ~20% of expressed genes and characterize the multiple lineage differentiation. Second, we profiled 16 HMs with ChIP-seq data available for all six cell types, including nine types of acetylation and seven types of methylation. Following the observation that the dynamic changes of most HMs are enriched in AS exons and significantly different between inclusion-gain and inclusion-loss exons, we found that 3 of the 16 investigated HMs (H3K36me3, H3K27ac, and H4K8ac) are strongly associated with 52.8% of hESC differentiation-related AS exons. We then linked the association between HMs and AS to cell-cycle progression based on the additional discovery that the AS genes predominantly function in cell-cycle progression. More intriguingly, we found that HMs and AS are associated in G2/M phases and involved in ESC fate decision through promoting pluripotency state dissolution, repressing self-renewal, or both. Especially, with experimental valuations, we demonstrated an H3K36me3-regulated isoform switch from PBX1a to PBX1b, which is implicated in hESC differentiation by attenuating the activity of the pluripotency regulatory network. Collectively, we presented a mechanism conveying the HM information into cell fate decision through the regulation of AS, which will drive extensive studies on the involvements of HMs in cell fate decision via determining the transcript structure rather than only the transcript abundance.

## Results

### 1. Alternative splicing characterizes hESC differentiation

The role of AS in the regulation of ES cell fates adds another notable regulatory layer to the know mechanisms that govern stemness and differentiation [57]. To screen the AS events associated with ES cell fate decision, we investigated a panel of RNA-seq data during hESC (H1 cell line) differentiation [16]. We considered four cell types directly differentiated from H1 cells, including trophoblast-like cells (TBL), mesendoderm (ME), neural progenitor cells (NPC), and mesenchymal stem cells (MSC). We also considered IMR90, a cell line for primary human fetal lung fibroblast, as an example of terminally differentiated cells. These cells represent five cell lineages of different developmental levels (Additional file 1: Figure S1A). We identified thousands of AS events of all types with their changes of 'per spliced in' (∆PSIs) are larger than 0.1 (inclusion-loss) or less than -0.1 (inclusion-gain), and with the FDRs are less than 0.05 based on the measurement used by rMATS [58] (Additional file 1: Figure S1B and Table S1, see Methods). We implemented further analyses only on the most common AS events, including 3,513 mutually exclusive exons (MXE) and 3,678 skipped exons (SE), which are referred to as hESC differentiation-associated AS exons (Additional file 1: Figure S1C and Additional file 2: Table S2).

These hESC differentiation-related AS exons possess typical properties as previously described [59, 60] as follows. (1) Most of their hosting genes are not differentially expressed between hESCs and differentiated cells (Additional file 1: Figure S1D). (2) They tend to be shorter with much longer flanking introns compared to the average length of all exons and introns (RefSeq annotation), respectively (Additional file 1: Figures S2A, B). (3) The arrangement of shorter AS exons surrounded by longer introns is consistent across cell lineages and AS types (Additional file 1: Figures S2C, D). (4) The lengths of AS exons are more often divisible by three to preserve the reading frame (Additional file 1: Figure S2E).

During hESC differentiation, about 20% of expressed genes undergo AS (2,257 genes for SE and 2,489 genes for MXE), including previously known ESC-specific AS genes, such as the pluripotency factor *FOXP1* [14] (Figure 1A) and the Wnt/*β*-catenin signalling component *CTNND1* [15] (Figure 1B). These hESC differentiation-related AS genes include many transcription factors, transcriptional co-factors, chromatin remodelling factors, housekeeping genes, and bivalent domain genes implicated in ESC pluripotency and development [41] (Figures 1C and Additional file 1: Figure S1C). Enrichment analysis based on a stemness gene set [61] also shows that hESC differentiation-related AS genes are enriched in the regulators or markers that are most significantly associated with stemness signatures of ESCs (Additional file 1: Figure S3A, see Methods).

**Figure 1.**
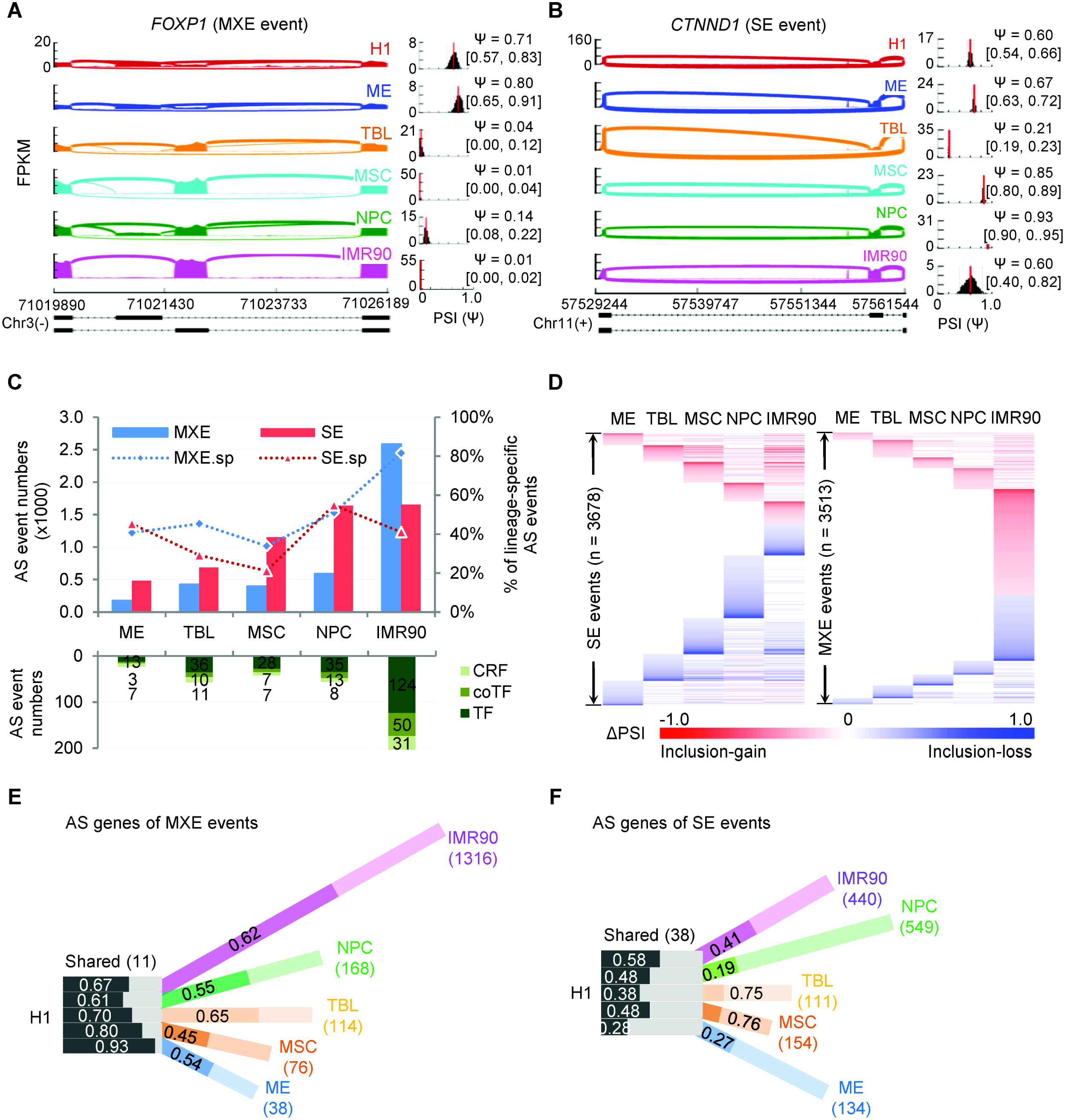
Alternative splicing characterizes the hESC differentiation. **(A)** and **(B)** Sashimi plots show two AS events of previously known ESC-specific AS events, *FOXP1* (A) and *CTNND1* (B). Inset histograms show the PSIs (ψ) of the AS exons in all cell types based on the MISO estimation. **(C)** The bar graph shows that the number of total AS events and lineage-specific AS events increase coordinately with the developmental levels. Higher developmental level induces more (lineage-specific) AS events. MXE.sp and SE.sp indicate the percentage of lineage-specific AS events. **(D)** Heat maps show the differential ' per cent splice in' (∆PSIs) of SE (left panel) and MXE (right panel) AS events (rows) for each cell lineage (columns). For MXE event, the ∆PSIs are of the upstream exons. **(E)** and **(F)** The hosting genes of MXE (E) and SE (F) AS events characterize cell lineages. Black-and-white bars refer to the common AS genes shared by all cell lineages, while the colour bars indicate the lineage-specific AS genes. The length of the colour bars is proportional to the percentage of lineage-specific genes. Dark fills indicate the inclusion-gain events, while light fills indicate the inclusion-loss events. The numbers in the bars are the proportion of corresponding parts; the numbers in the parentheses are the numbers of common AS genes or lineage-specific AS genes of each lineage. Gain or loss for MXE events refers to the upstream exons. Also, see **Figures S1-S3**.

Clustering on AS events across cell lineages show lineage-dependent splicing patterns (Figure 1D). Upon hESC differentiation, the SE exons tend to lose their inclusion levels (inclusion-loss), while the upstream exons of MXE events are likely to gain their inclusion levels (inclusion-gain) (Fisher's exact test, p = 3.83E-107). The numbers of AS events increase accordingly with the developmental level following hESC differentiation (Figure 1C). For example, the differentiation to ME involves the fewest AS events and ME presents the most stem-cell-like AS profiles, while the IMR90 has the most AS events and exhibits the most similar AS profiles to adult cells (Figure 1C, D). Inter-lineage comparisons show, on average, that 42.0% of SE and 56.4% of MXE events (Figures 1C, D and Additional file 1: Figure S3B, C), involved in 29.6% and 38.6% of AS hosting genes (Figures 1E, F and Additional file 1: Figure S3D, E), are lineage-specific. In contrast, only 0.65% of SE and 0.14% of MEX events (Additional file 1: Figures S3B, C), involved in 0.49% and 1.52% of AS hosting genes, are shared by all lineages (Figures 1E, F and Additional file 1: Figure S3D, E). Similar trends are observed from pairwise comparisons (Additional file 1: Figure S3F). Furthermore, one-third of AS genes (n = 881) have both MXE and SE events (Additional file 1: Figure S3G). Only four genes are common across all cell lineages and AS types, of which the AS events of *Ctnnd1* and *Mbd1* have been reported to regulate mESC differentiation [15]. Together, these results demonstrate that alternative splicing depicts lineage- and developmental level-dependent characterizations of hESC differentiation.

### 2. Dynamic changes of histone modifications predominantly occur in AS exons

In ESCs, epigenetic mechanisms contribute mainly to maintaining the expression of pluripotency genes and the repression of lineage-specific genes in order to avoid exiting from stemness. Upon differentiation, epigenetic mechanisms orchestrate the expression of developmental programs spatiotemporally to ensure the heritability of existing or newly acquired phenotypic states. Though epigenetic signatures are mainly found to be enriched in promoters and enhancers, it has become increasingly clear that they are also present in gene bodies, especially in exon regions, implying a potential link of epigenetic regulation to pre-mRNA splicing [62, 63]. Consistent with previous reports [37, 38, 64], we also observed that few involved splicing factors are differentially expressed during H1 cells differentiation (Additional file 1: Figure S3H, see Methods), which confirms the existence of an additional layer of epigenetic regulations on AS. However, the extents to which the AS is epigenetically regulated and how these AS genes contribute to the cell fate decision are poorly understood. We focused on 16 histone modifications, including nine histone acetylation and seven histone methylation that have available data in all six cell types (see Methods), and aimed to reveal their associations with AS genes during hESC differentiation.

To investigate whether the dynamic changes of these HMs upon cell differentiation prefer to AS exons consistently (Figures 2A, B), we profiled the differential HM patterns of around the hESC differentiation-associated AS exons and the same number of randomly selected constitutive splicing (CS) exons of the same AS genes for each differentiation lineage. We compared the changes of ChIP-seq reads count (normalized ∆s reads count, see Methods) in ±150bp regions around the splice sites upon hESC differentiation (Figure 2C and Additional file 1: Figure S4, see Methods). Except for a small part of cases (with black dots or boxes in Figure 2D), most HMs changed more significantly around AS exons than around constitutive exons upon hESC differentiation (Mann-Whitney-Wilcoxon test, p ≤ 0.05, Figures 2D and Additional file 1: Figure S4). Nevertheless, some HMs displayed strong links to AS, such as H3K79me1 and H3K36me3, while others only had weak link strengths, such as H3K27me3 and H3K9me3 (Figure 2D). This result is consistent with the fact that the former are involved in active expression and AS regulation [39, 46, 65], while the latter are the epigenetic marks of repressed regions and heterochromatin [66]. The link strengths are presented as the -log10 p-values to test whether the HM changes consistently prefer the AS exons across different cell lineages and AS types (Figure 2D sidebar graph, see Methods). Taken together, these results, from a global view, revealed a potential regulatory link from HMs to RNA splicing, of which some are strong while the others are weak.

**Figure 2.**
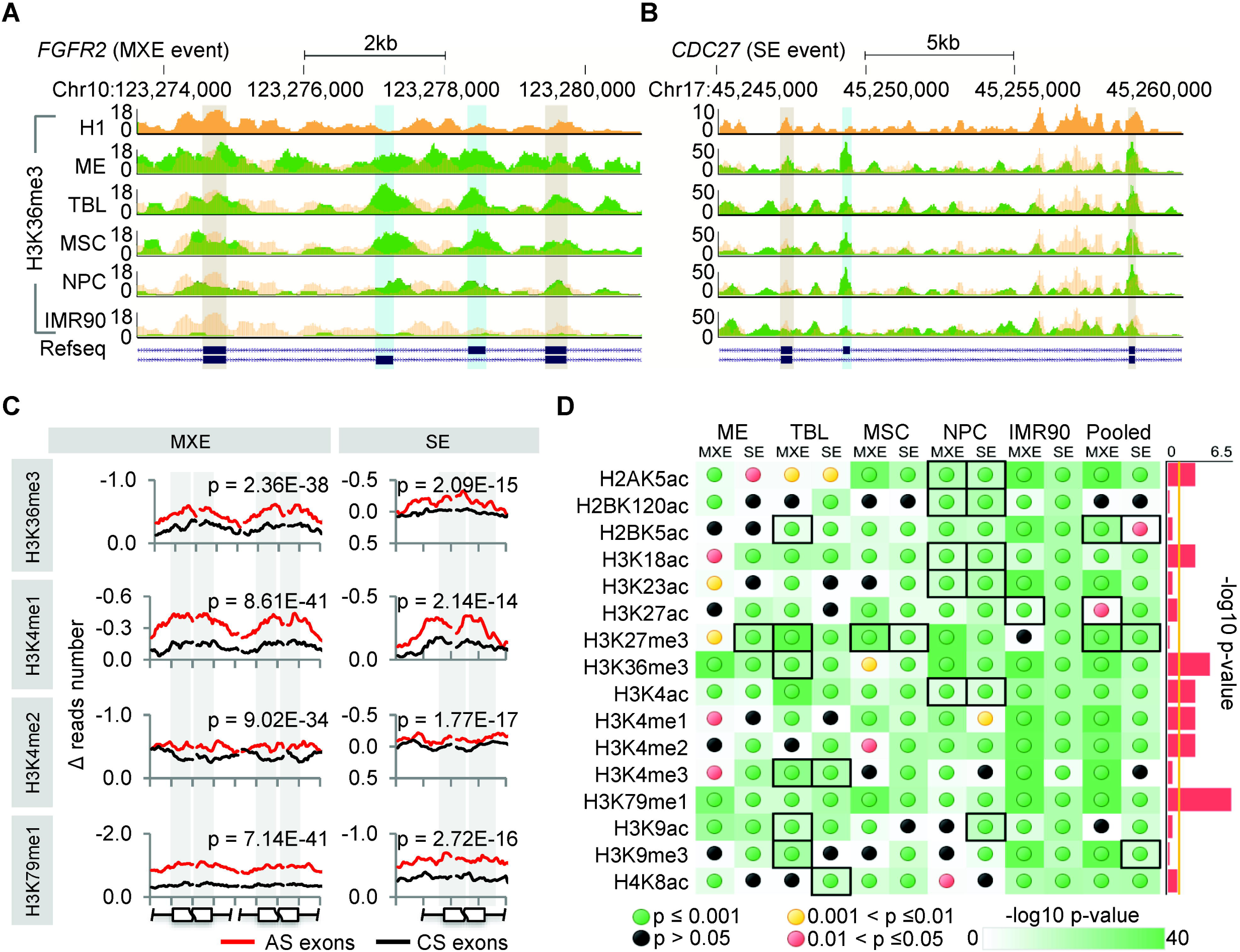
Dynamic changes of histone modifications predominantly occur in AS exons. **(A)** and **(B)** Genome browser views of representative H3K36me3 changes in MXE (exemplified as *FGFR2)* and SE (exemplified as *CDC27)* events respectively, showing that the changes of H3K36me3 around the AS exons (blue shading) are more significant than around the flanking constitutive exons (gray shading) in 4 H1-derived cell types and IMR90. The tracks of H1 are duplicated as yellow shadings overlapping with other tracks of the derived cells (green) for a better comparison. **(C)** Representative profiles of HM changes (normalized ∆ reads number) around the AS exons and randomly selected constitutive splicing (CS) exons upon hESC differentiation, shown as the average of all cell lineages pooled together. The ±150bp regions (exons and flanking introns) of the splice sites were considered and 15bp-binned to produce the curves. It shows that the changes of HMs are more significant around AS exons than around constitutive exons, especially in exonic regions (grey shading). The p-values, Mann-Whitney-Wilcoxon test. (D) The statistic significances for changes of all 16 HMs in all cell lineages and pooling them together (pooled), represented as the -log10 p-values based on Mann-Whitney-Wilcoxon test. The detailed profiles are provided in **Figure S4**. Black boxes indicate the cases that HMs around constitutive exons change more significantly than around AS exons, corresponding to the red-shaded panels in **Figure S4**. Sidebars represent the significances whether the changes of HMs are consistently enriched in AS exons across cell lineages, showing the link strength between AS and HMs and represented as the - log10 p-value based on Fisher's exact test. The yellow vertical line indicates the significance cutoff of 0.05. Also, see **Figure S4**.

### 3. Three histone modifications are significantly associated with alternative splicing upon hESC differentiation

To quantitatively associate the HMs with AS, all ChIP-seq data were processed for narrow peak calling using MACS2 [67]. For each AS exon of each differentiation lineage, we then quantified the differential inclusion levels, i.e., the changes of 'percent splice in' (∆PSIs, Additional file 1: Figure S1B), and the differential HMs signals, i.e., the changes of normalized narrow peak height of ChIP-seq (∆HMs, Additional file 1: Figure S5A, see Methods) between H1 and differentiated cells. We observed significant differences in all HM profiles (except H3K27me3, Additional file 1: Figure S5B) between the inclusion-gain and inclusion-loss exons across cell lineages and AS types (Mann-Whitney-Wilcoxon test, p ≤ 0.05) (Figure 3A and Additional file 1: Figure S5B). However, three independent correlation tests showed only weak global quantitative associations between the ∆PSIs and ∆HMs for some HMs (Figure 3C and Additional file 1: Figure S5C), including eight HMs for MXE AS exons and eight HMs for SE AS exons. The weak associations may indicate that only subsets of AS exons are strongly associated with HMs, and *vice versa*, which is consistent with a recent report [68].

**Figure 3.**
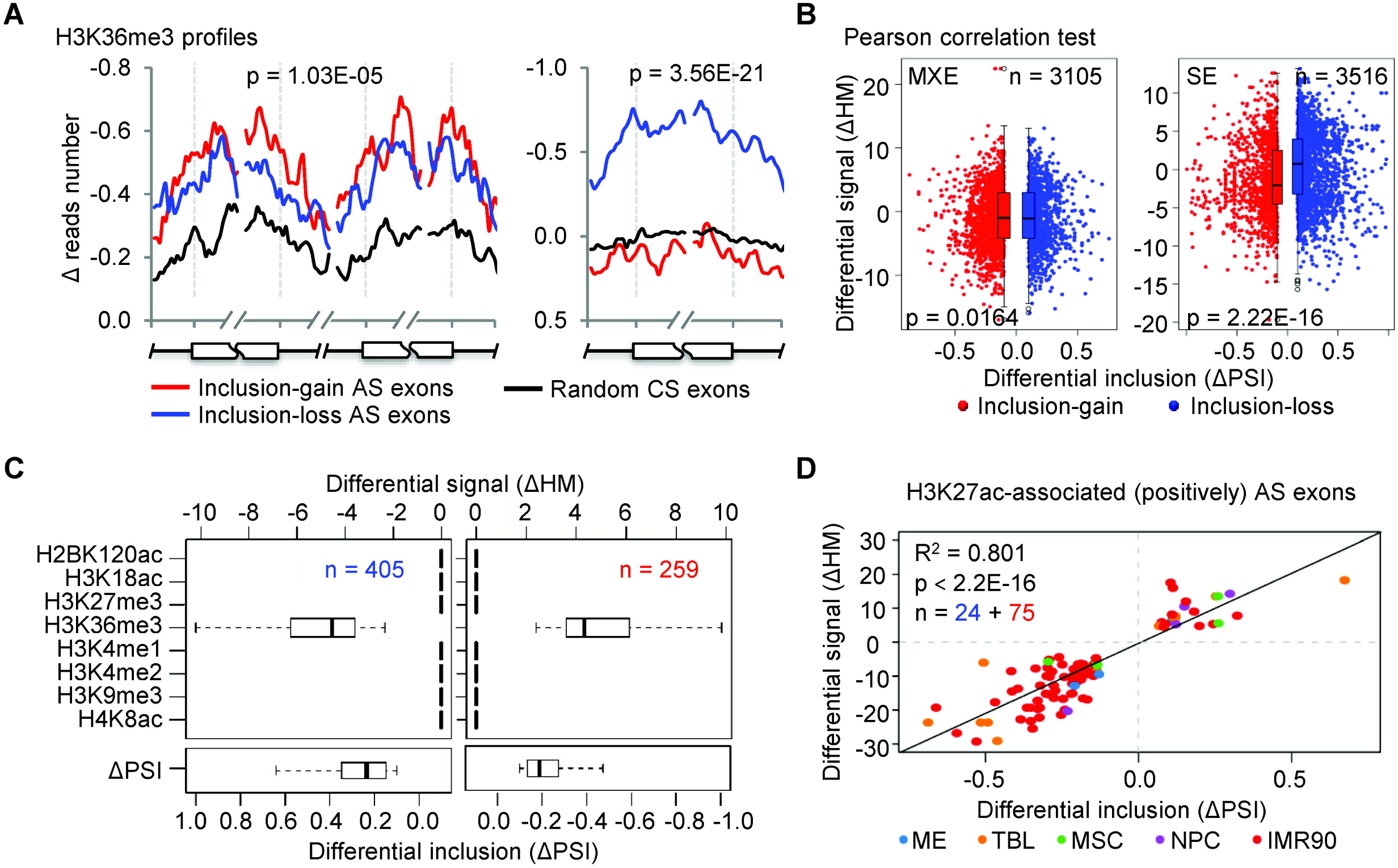
A subset of histone modifications and alternative splicing are strongly associated upon hESC differentiation. **(A)** Representative profiles of HM (H3K36me3) changes (normalized ∆ reads number) around the inclusion-gain (red lines) and inclusion-loss (blue lines) AS exons, as well as randomly selected constitutive splicing (CS) exons (black lines) for both MXE (left panel) and SE (right panel) AS events. It shows that HM changes are significantly different between inclusion-gain and inclusion-loss AS exons (p-values, Mann-Whitney-Wilcoxon test). **Figure S5B** provides the whole significances of all HMs across AS types and cell lineages. **(B)** Pearson correlation test between differential HM signals (∆HMs) and differential inclusion levels (∆PSIs), taking H3k36me3 as an example. **Figure S5C** provides the correlation test results of other HMs based on two more tests. **(C)** A representative k-means cluster shows a subset of SE AS events having a negative correlation between the ∆PSIs and the ∆HMs of H3K36me3. **Figures S5D** and **S6** provide all the clustering results. **(D)** Scatter plot shows that HM-associated AS events display significant correlations between the ∆PSIs and the ∆HMs upon hESC differentiation, taking H3K27ac–associated (positively) MXE events as an example. Also, see **Figures S5-S6**.

To explore the subsets of highly associated AS exons and corresponding HMs, we performed *k*-means clustering on the sets of inclusion-gain and inclusion-loss exons of SE and MXE events, separately, taking the ∆HMs of eight identified HMs as epigenetic features (Figure 3C and Additional file 1: Figure S5D and S6, see Methods). We obtained three subsets of HM-associated SE exons and three subsets of HM-associated MXE exons (Additional file 3: Table S3). The three HM-associated SE subsets include 180, 664, and 1,062 exons, and are negatively associated with H4K8ac (Additional file 1: Figure S6), negatively associated with H3K36me3 (Figure 3C), and positively associated with H3K36me3 (Additional file 1: Figure S6), respectively. The three HM-associated MXE subsets include 99, 821, and 971 exons, and are positively associated with H3K27ac (Figure 3D), negatively associated with H3K36me3 (Additional file 1: Figure S6), and positively associated with H3K36me3 (Additional file 1: Figure S6), respectively. The exons of each subset show significant correlations between their ∆PSIs and ∆HMs upon hESC differentiation (Figure 3D). These HM-associated AS exons account for an average of 52.8% of hESC differentiation-related AS events, on average (Additional file 1: Figure S5E).

Of the three AS-associated HMs, H3K36me3 has both positive and negative correlations with AS exons. This is consistent with the fact that H3K36me3 has dual functions in regulating AS through two different chromatin-adapter systems, PSIP1/SRSF1 [47] and MRG15/PTBP1 [46]. The former increases the inclusion levels of targeting AS exons, whereas the latter decreases the inclusion levels [39]. As expected, 139 and 11 of our identified H3K36me3-associated AS genes have been reported to be regulated by SRSF1 [69, 70] (Additional file 1: Figure S5F) and PTBP1 [71] (Additional file 1: Figure S5G), respectively. Taken together, our analysis showed that more than half (52.8%) of hESC differentiation-associated AS events are significantly associated with 3 of 16 HMs during hESC differentiation, including H3K36me3, H3K27ac, and H4K8ac.

### 4. HM-associated AS genes predominantly function in G2/M phases to facilitate hESC differentiation

Epigenetic mechanisms have been proposed to be dynamic and play crucial roles in human ESC differentiation [16, 17]. Given the aforementioned associations between HMs and AS, and the well-established links between AS and hESC differentiation, we hypothesized that the three HMs (H3K36me3, H3K27ac, and H4K8ac) may contribute to stem cell differentiation through their associated AS events. To test our hypothesis and gain more insights into the differences between the HM-associated and - unassociated AS events, we performed comparative function analyses between their hosting genes, revealing that HMs are involved in alternatively splicing the core components of cell-cycle machinery and related pathways to regulate stem cell pluripotency and differentiation.

We found that HMs prefer to be associated with even shorter AS exons (Additional file 1: Figure S7A, p < 0.001, Student's t-test), though AS exons are shorter than the average length of all exons (Additional file 1: Figure S2A). HM-associated genes (n = 2125) show more lineage specificity, i.e., more genes (49.76% versus 29.6% of MXE or 38.6% of SE genes) are lineage-specific (Additional file 1: Figures S7B and S3D, E), regardless of whether IMR90 is included or not (Figure S7C). Only a few HM-associated genes are shared by different cell lineages, even in pairwise comparisons (Additional file 1: Figure S7D); the most common shared genes are lineage-independent housekeeping genes (Additional file 1: Figure S7E). These suggest that HM-associated AS genes contribute more to lineage specificity. In addition, the HM-associated AS genes (966 of 2,125) are more enriched in stemness signatures than do unassociated AS genes (429 of 1,057) (Figure 4A). TF binding enrichment analysis shows that HM-associated AS genes are likely to be regulated by TFs involved in cell differentiation, whereas HM-unassociated AS genes are likely to be regulated by TFs involved in cell proliferation and growth (Figure 4B). All these results suggest that HM-associated and -unassociated AS genes function differently during hESC differentiation. Gene ontology (GO) enrichment analysis shows that more than half of the HM-associated AS genes (1,120 of 2,125) function in cell-cycle progression and exhibit more significant enrichment than do HM-unassociated AS genes (376 of 1,057, Figures 4C, D and Additional file 1: Figure S8A). The significance of the top enriched GO term (GO:0007049, cell cycle) is consistent across cell lineages, although HM-associated AS genes exhibit more lineage specificity and few of them are shared among lineages (Additional file 1: Figures S7B-D and S8B). These results suggest the involvement of HMs in AS regulation of the cell-cycle machinery that has been reported to be exploited by stem cells to control their cell fate decision [21].

**Figure 4.**
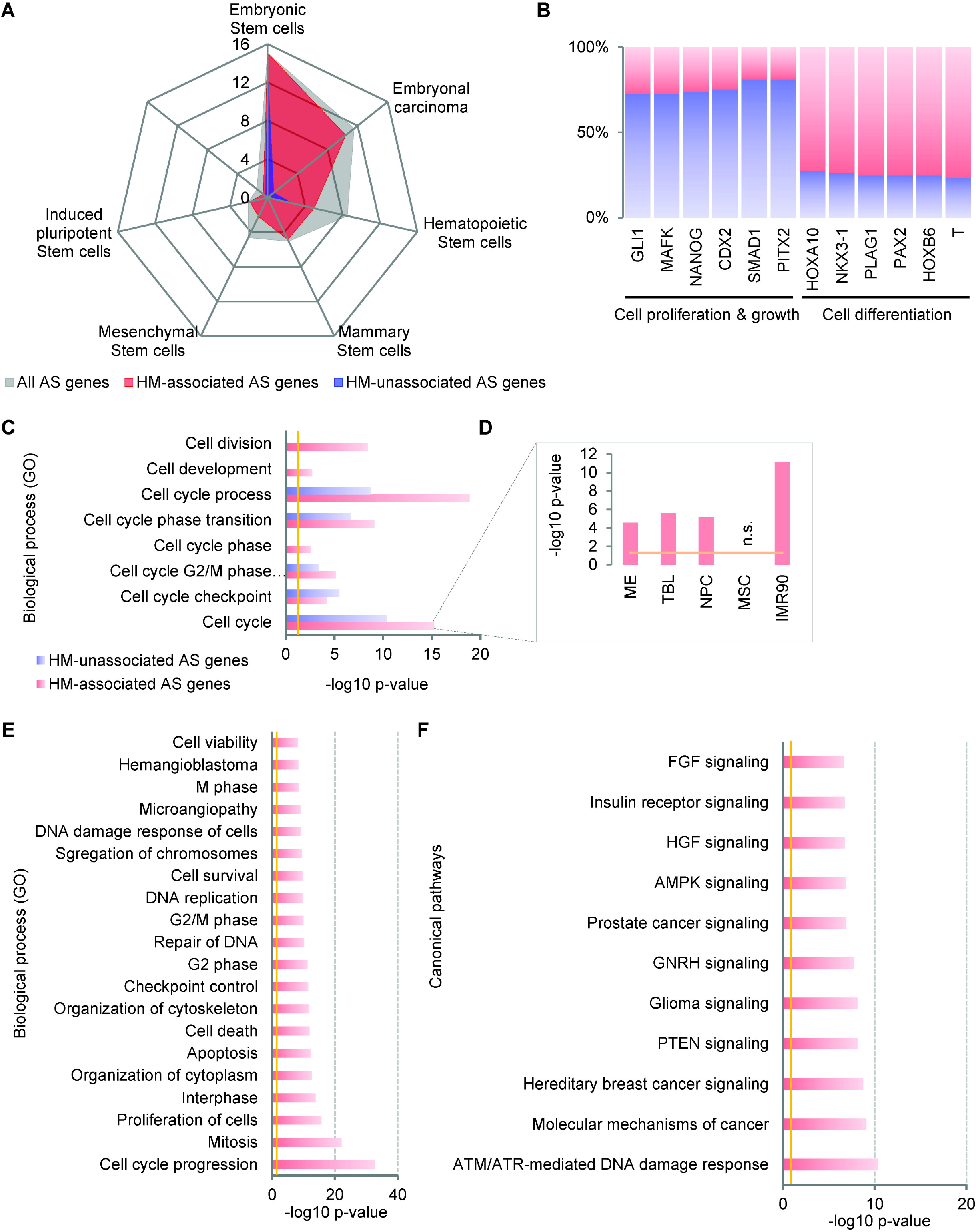
HM-associated AS genes predominantly function in G2/M cell-cycle phases contributing to hESC differentiation. **(A)** HM-associated AS genes are enriched more significantly in stemness signatures than HM-unassociated AS genes. **(B)** TF binding enrichment shows that HM-associated AS genes prefer to be regulated by TFs involved in cell differentiation, while the HM-unassociated AS genes are prone to be regulated by TFs involved in cell proliferation and growth. **(C)** GO enrichment analysis shows that HM-associated AS genes are enriched more significantly in cell cycle progression than HM-unassociated AS genes, shown as the -log10 p-values after FDR (≤ 0.05) adjustment. **(D)** The significant enrichment of HM-associated AS genes in the cell cycle are consistent across cell lineages, with the MSC as an exception that no significant enrichment was observed. **(E)** The top twenty enriched functions show that HM-associated AS genes involved in cell cycle progression prefer to function in G2/M phases and DNA damage response. **(F)** The canonical pathway enrichment shows that AMT/ATR-mediated DNA damage response is the top enriched pathway of HM-associated AS genes. The vertical lines (yellow) indicate the significance cutoff of **0.05**. Also, see **Figures S7-S8**.

Further study of the top enriched cell-cycle AS genes (Figure 4D and Additional file 1: Figure S8A) shows that HM-associated (n = 282) and -unassociated AS genes (n = 150) play roles in different cell-cycle phases and related pathways. The former is prone to function in G2/M phases and DNA damage response (Figures 4E, F). This indicates that HMs contribute to cell differentiation, at least partially, via AS regulations in these phases, which is consistent with the fact that inheritance of HMs in daughter cells occurs during the G2 phases [21]. The latter play roles in G1 phase, cell cycle arrest, and Wnt/|3-catenin signalling (Additional file 1: Figures S8C, D). Since cell fate choices seem to occur or at least be initiated during G1/S transition [55], while cell fate commitment is achieved in G2/M [56], it could be rational for stem cells to change their identity during the G2 phase when HMs are reprogrammed [21].

Intriguingly, the top enriched pathway of HM-associated AS genes is 'ATM/ATR-mediated DNA damage response', which is activated in S/G2 phases and has been recently reported as a gatekeeper of the pluripotency state dissolution (PSD) that participates in allowing hESC differentiation [56]. Together with our previous results [20], it suggests the presence of a combinational mechanism involving HMs and AS, wherein HMs facilitate the PSD and cell fate commitment by alternatively splicing the key components of the ATM/ATR pathway. Additionally, many cell-cycle TF genes are involved in the top enriched HM-associated AS gene set. The pre-B-cell leukaemia transcription factor 1 (*PBX1*) is one of these genes that contribute to cell cycle progression and is discussed later in next section. Taken together, we suggest that 3 of 16 HMs function in positive or negative ways affect the AS of subsets of genes, and further contribute to hESC differentiation in a cell-cycle phase-dependent manner. The results suggest a potential mechanistic model connecting the HMs, AS regulations, and cell-cycle progression with the cell fate decision.

### 5. Splicing of PBX1 links H3K36me3 to hESC fate decision

The past few years have identified key factors required for maintaining the pluripotent state of hESCs [72, 73], including NANOG, OCT4 (POU5F1), SOX2, KLF4, and c-MYC, the combination of which was called Yamanaka factors and sufficient to reprogram somatic cells into induced pluripotent stem cells (iPSCs) [74]. These factors appear to activate a transcriptional network that endows cells with pluripotency [75]. The above integrative analyses showed strong links between three HMs and RNA splicing, revealing a group of epigenetic regulated AS genes involved in cell cycle machinery. *PBX1* was one of the genes that their ASs are positively associated with H3K36me3 (Figures 5A, B). Its protein is a member of the TALE (three-amino acid loop extension) family homeodomain transcription factors [76, 77] and well known for its functions in lymphoblastic leukaemia [78-81] and several cancers [82-91]. PBX1 also plays roles in regulating developmental gene expression [92], maintaining stemness and self-renewal [82, 93, 94], and promoting the cell-cycle transition to S phase [95]. Additionally, multiple lines of evidence obtained from *in vivo* and in vitro highlighted its functions as a pioneer factor [88, 96]. However, few studies have distinguished the distinct functions of its different isoforms.

**Figure 5.**
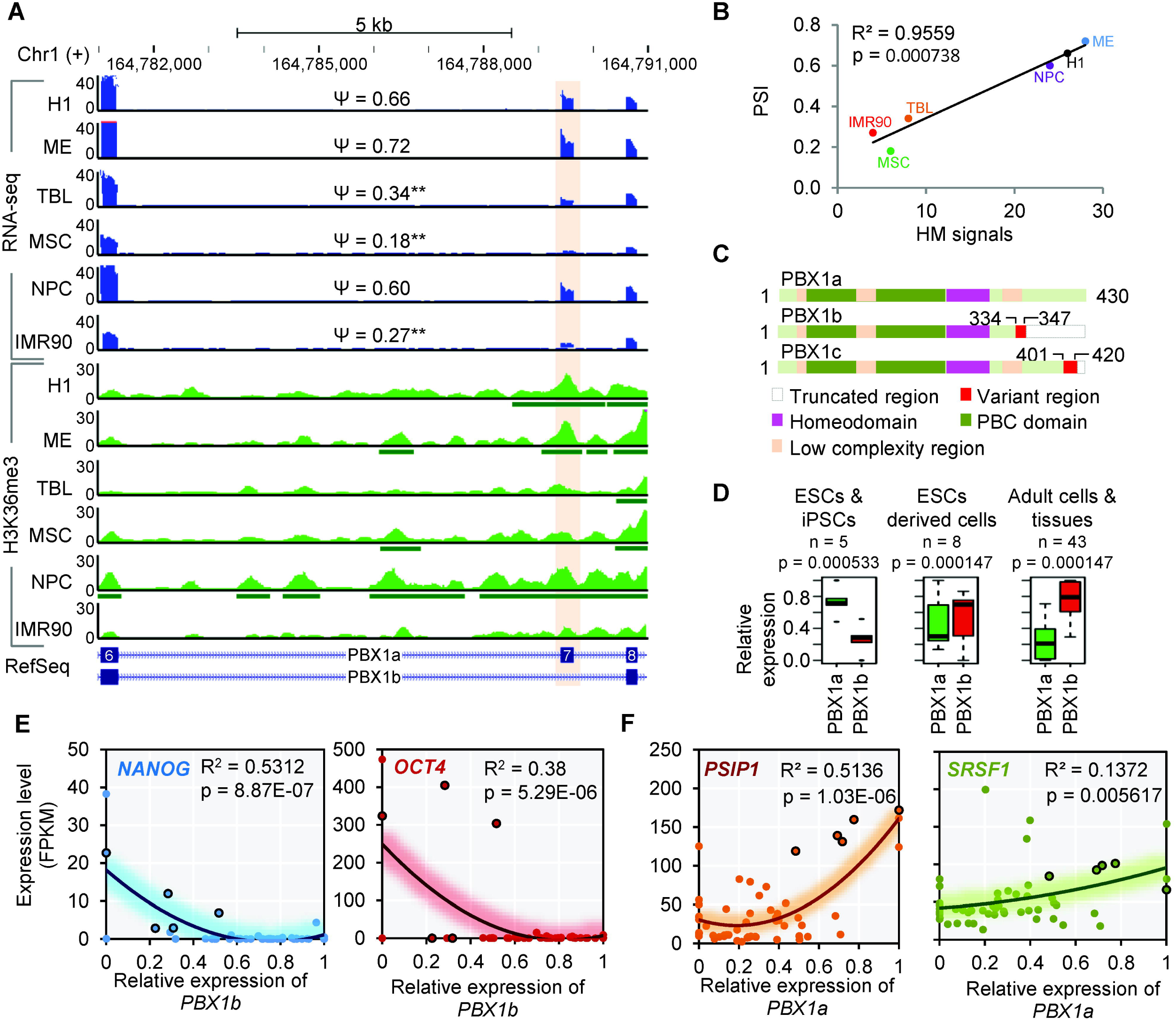
Isoform switch from PBX1a and PBX1b during hESC differentiation. **(A)** Genome browser view shows the AS event and H3K36me3 signals of PBX1 upon hESC differentiation. The green horizontal bars below the ChIP-seq tracks indicate the narrow peaks called by MACS2. **(B)** The inclusion level for exon 7 of PBX1 is significantly correlated to the H3K36me3 signals over this exon across cell lineages. **(C)** The sequence difference of three protein isoforms of PBX1 and the main functional domains. **(D)** The relative expressions of PBX1a and PBX1b in 56 cells/tissues, representing the differential expressions of two isoforms in three groups based on their developmental states. **(E)** The expression levels of NANOG and OCT4 genes are negatively correlated with the expression of PBX1b. **(F)** The expression levels of PSIP1 and SRSF1 show significant positive correlations with the expression level of PBX1a. Also, see **Figure S9-S10**.

PBX1 has three isoforms [97], including PBX1a, PBX1b, and PBX1c (Figure 5C and Additional file 1: Figure S9A). PBX1a and PBX1b are produced by the alternative splicing of exon 7 (Figure 5A) and attract most of the research attention of PBX1. PBX1b retains the same DNA-binding homeodomain as PBX1a, but changes 14 amino acids (from 334 to 347) and truncates the remaining 83 amino acids at the C-terminus of PBX1a (Figure 5C and Additional file 1: Figure S9A). This C-terminal alteration of PBX1a has been reported to affect its cooperative interactions with HOX partners [98], which may impart different functions to these two isoforms. We here revealed its H3K36me3-regulated isoform switch between PBX1a and PBX1b, which functions at the upstream of pluripotency transcriptional network to link H3K36me3 with ESC fate decision.

We first observed differential transcript expressions of these two isoforms between the hESCs and differentiated cells, wherein *PBX1a* was predominantly transcribed in hESCs, while *PBX1b* was predominantly induced in differentiated cells (Figure 5A and Additional file 1: Figure S9B). The same trend was also observed in an extended dataset of 56 human cell lines/tissues (Figure 5D) from the Roadmap [99] and ENCODE [100] projects (Additional file 4: Table S4). Additionally, we did not observe significantly different expression of the total *PBX1* and three other *PBX* family members across cell types (Additional file 1: Figure S9C, fold change < 2), indicating that the isoform switch of *PBX1*, rather than the differential expression of its family members play more important roles during hESC differentiation. To further test the possible mechanism by which PBX1b contributes to stem cell differentiation, we investigated the transcription levels of Yamanaka factors. Of these TFs, the *NANOG* is activated by direct promoter binding of PBX1 and KLF4, which is essential for stemness maintenance [93, 101]. Consistently, all these core factors are repressed in differentiated cells where *PBX1b* is highly expressed (Additional file 1: Figures S9D-G), even though the *PBX1a* is expressed. Based on the 56 human cell lines/tissues, we also observed significant negative correlations between expression of most important pluripotent factors *(NANOG* and *OCT4)* and *PBX1b* (Figure 5E), as well as positive correlations between these two factors and *PBX1a* (or inclusion level of exon 7, Additional file 1: Figures S10A-B). Consistent with previous reports showing that the PBX1a and PBX1b differ in their ability to activate or repress the expression of reporter genes [102, 103], we hypothesize that PBX1a promotes the activity of the pluripotent regulatory network by promoting the expression of *NANOG*, whereas PBX1b may attenuate this activity by competitively binding and regulating the same target gene, since PBX1b retains the same DNA-binding domain as PBX1a. These observations are strongly suggestive that the switch from PBX1a to PBX1b is a mechanism by which PBX1 contributes to hESC differentiation via regulating the pluripotency regulatory network.

The exon 7 of PBX1 shows significantly positive correlations between its inclusion levels (PSIs) and the surrounding epigenetic signals of H3K36me3 in hESCs and differentiated cells (Figure 5B). It suggests a potential role of H3K36me3 in regulating the isoform switch between *PBX1a* and *PBX1b*. To investigate the regulatory role of H3K36me3, we focused on two previously proved chromatin-adaptor complexes, MRG15/PTBP1 [46] and PSIP1/SRSF1 [47], which endow dual functions to H3K36me3 in AS regulation [39]. Based on the 56 cell lines/tissues from the Roadmap/ENCODE projects, we first found significant positive correlations between the expressions of *PBX1a* (or inclusion level of exon 7) and *PSIP1/SRSF1* (Figure 5F), but not with *MRG15/PTBP1* (Additional file 1: Figures S10C-D). This result suggests that the alternative splicing of *PBX1* is epigenetic regulated by H3K36me3 through the PSIP1/SRSF1 adaptor system, which was strongly supported by a recent report using the HeLa cell lines [69]. The overexpression of SRSF1 in Hela cells introduces a PSI increase of 0.18 for the exon 7 of PBX1 (chr1: 164789308-164789421 based on NCBI37/hg19 genome assembly) based on the RNA-seq (Table S1 of [69]). Additionally, this exon was one of the 104 AS exons that were further validated using radioactive RT-PCR (Table S2 of [69]). Their results showed that exon 7 of PBX1 is indeed a splicing target of SRSF1, supporting our conclusions.

We then validated the above hypotheses on MSCs and IM90 cells, since these two cells types show the most significant difference from H1 cells regarding our hypotheses (Figure 5B). We cultured H1 cells, IMR90 cells, and induced H1 cells to differentiate into MSCs (H1-MSCs, see Methods for details). Additionally, we also included other two sources of MSCs, including one derived from human bone marrow (hBM-MSCs) and the other derived from adipose tissue (hAT-MSCs) (see Methods for details). Consistent with the results from RNA-seq, the same expression patterns of Yamanaka factors in H1, MSCs, and IMR90 cells were observed using qRT-PCR and Western blot (Figure 6A), which confirmed the pluripotent state of H1 cells and the differentiated states of other cell types. We then detected the isoform switch from PBX1a to PBX1b in our cultured cells, which are consistent both in mRNA and protein levels (Figure 6B and Additional file 1: Figure S10E) and further confirmed by the western blot using PBX1b-specific antibody (anti-PBX1b) (Figure 6B bottom and Additional file 1: Figure S10E iii). These results have verified that the PBX1b was significantly induced in differentiated cells, where the PBX1a was significantly reduced.

**Figure 6.**
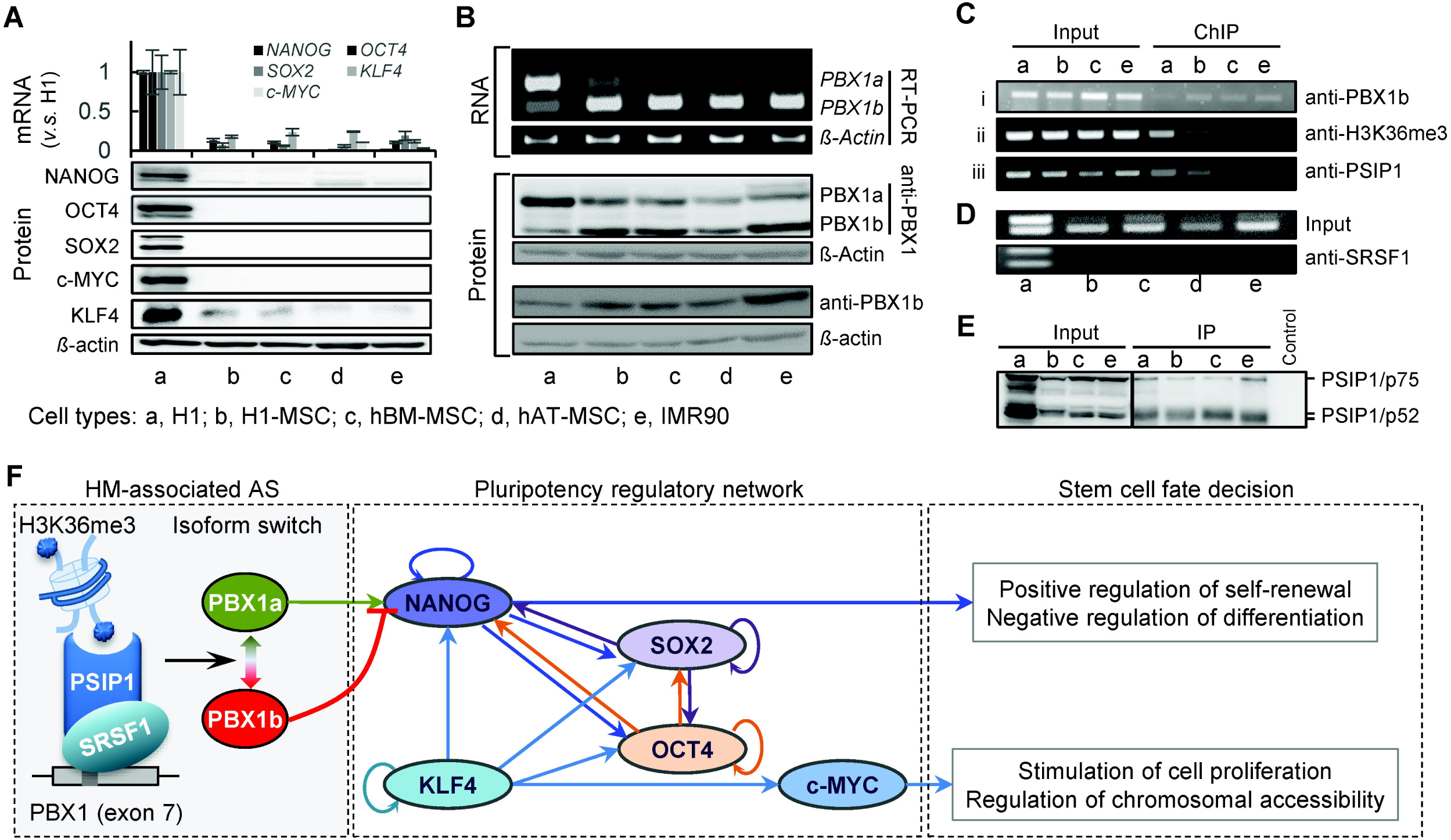
Isoform switch of PBX1 links H3K36me3 to hESC fate decision. **(A)** qRT-PCR and western blot show the expression levels of Yamanaka factors in H1, MSC and IMR90 cells. Whiskers denote the standard deviations of three replicates. **(B)** RT-PCR and western blot show the isoform switches between PBX1a and PBX1b from H1 cells to differentiated cells. **(C) i**. ChIP-PCR shows the differential binding of PBX1b to NANOG promoter in H1 cells and differentiated cells; **ii**. ChIP-PCR shows the reduced H3K36me3 signal in differentiated cells; iii. ChIP-PCR shows the differential recruitment of PSIP1 to the exon **7** of PBX1. **(D)** RIP-PCR show the differential recruitment of SRSF1 around the exon **7** of PBX1. **(E)** Co-IP shows the overall physical interaction between PSIP1 and SRSF1 in all studied cell types. **(F)** The mechanism by which H3K36me3 is linked to cell fate decision by regulating the isoform switch of PBX1, which functions upstream of the pluripotency regulatory network. Also, see **Figures S9-S10**.

We also validated the mechanism by which the splicing of PBX1 links H3K36me3 to stem cell fate decision. We first confirmed that PBX1b also binds to the promoter of *NANOG* at the same region where PBX1a binds to, and the binding signals (ChIP-PCR) were high in the differentiated cells but very low in H1 stem cells (Figure 6C i and Additional file 1: Figure S10F i). Consistent with the results from ChIP-seq, we also observed reduced H3K36 tri-methylation around the exon 7 of PBX1 based on ChIP-PCR assay (Figure 6C ii and Additional file 1: Figure S10F ii). Furthermore, the chromatin factor PSIP1 only showed high binding signal in H1 stem cells (Figure 6C iii and Additional file 1: Figure S10F iii), which recruit the splicing factor SRSF1 to the PBX1 exclusively in H1 stem cells (Figure 6D and Additional file 1: Figure S10G) even though the physical binding between these two factors were universally detected in all cell types (Figure 6E). All these experimental results suggested that, upon differentiation, stem cells reduced the H3K36 tri-methylation and may attenuate the recruitment of PSIP1/SRSF1 adaptor around the exon 7 of PBX1, leading to the exclusion of exon 7 and highly expressed PBX1b in differentiated cells. High expression of PBX1b may attenuate the activity of PBX1a in promoting the pluripotency regulatory network.

Taken together, we suggested that H3K36me3 regulates the alternative splicing of PBX1 via the PSIP1/SRSF1 adaptor system, leading the isoform switch from PBX1a to PBX1b during hESC differentiation. Subsequently, PBX1b competitively binds to *NANOG* and abolishes the bindings of PBX1a. This competitive binding attenuates the pluripotency regulatory network to repress self-renewal and consequently facilitate differentiation (Figure 6F). These findings revealed how the PBX1 contributes to cell fate decision and exemplify the mechanism by which alternative splicing links histone modifications to stem cell fate decision.

## Discussion

ESCs provide a vital tool for studying the regulation of early embryonic development and cell-fate decision.[1]. In addition to the core pluripotency regulatory network, emerging evidence revealed other processes regulating ESC pluripotency and differentiation, including histone modifications (HMs), alternative splicing (AS), cell-cycle machinery, and signalling pathways [56]. Here, we connected these previously separate avenues of investigations, beginning with the discovery that 3 of 16 HMs are significantly associated with more than half of AS events upon hESC differentiation. Further analyses implicated the association of HMs, AS regulation, and cell-cycle progression with hESC fate decision. Specifically, HMs orchestrate a subset of AS outcomes that play critical roles in cell-cycle progression via the related pathways, such as ATM/ATR-mediated DNA response [20], and TFs, such as PBX1 (Additional file 1: Figure S10H). In this way, HMs, AS regulation, and signalling pathways are converged into the cell-cycle machinery that has been claimed to rule the pluripotency [21].

Although epigenetic signatures, such as HMs, are mainly enriched in promoters and other intergenic regions, it has become increasingly clear that they are also present in the gene body, especially in exon regions. This indicates a potential link between epigenetic regulation and the predominantly co-transcriptional occurrence of AS. Thus far, H3K36me3 [46, 47], H3K4me3 [44], H3K9me3 [45], and the acétylation of H3 and H4 [48-52] have been revealed to regulate AS, either via the chromatin-adapter systems or by altering Pol II elongation rate. Here, we investigated the extent to which the HMs could associate with AS by integrative analyses on both transcriptome and epigenome data during hESC differentiation. We found that three HMs are significantly associated with about half of AS events. By contrast, a recent report showed that only about 4% of differentially regulated exons among five human cell lines are positively associated with three promoter-like epigenetic signatures, including H3K9ac, H3K27ac and H3K4m3 [68]. Like that report, we also found a positive association of H3K27ac with a subset of AS events. However, our results differ regarding the other two HMs that we identified to be associated with AS.

In our study, H3K36me3 is associated with the most identified HM-associated AS events, either positively or negatively. It is reasonable since H3K36me3 is a mark for actively expressed genomes [65], and it has been reported to have dual roles in AS regulations through two different chromatin-adapter systems, PSIP1/SRSF1 [46] and MRG15/PTBP1 [47]. SRSF1 is a splicing factor which will increase the inclusion of targeted AS exons, and PTBP1 will decrease the inclusion levels of the regulated AS exons. Therefore, the exons that are regulated by the PSIP1/SRSF1 adapter system will show positive correlations with the H3K36me3, while the exons regulated by MRG15/PTBP1 will show negative correlations. Our results are consistent with this fact and show both direction correlations between different sets of AS events and H3K36me3. Many of these AS events in our study have been validated by other studies (Additional file 1: Figures S5F-G).

H4K8ac is associated with the fewest number of AS events in our results. Although rarely studied, H4K8ac is known to act as a transcriptional activator in both the promoters and transcribed regions [104]. Its negative association with AS is supported by the finding that it recruits proteins involved in increasing the Pol II elongation rate [105]. This suggests that H4K8ac may function in AS regulation by altering the Pol II elongation rate, rather than via the chromatin-adaptor systems. However, further studies are required. Collectively, possible reasons for the different results between this study and others [68] could be that (1) we considered differentiation from hESCs to five different cell types, which covered more inter-lineage AS events than the previous report; or (2) different sets of epigenetic signatures were considered, which may lead to relatively biased results in both studies. Obviously, the inclusion of more cell lineages and epigenetic signatures may reduce this bias. Therefore, an extended dataset of 56 cell lines/tissues were included in our study and the observations support our results.

Our study also extended the understanding that HMs contribute to cell fate decision via determining not only what parts of the genome are expressed, but also how they are spliced [39]. We demonstrated that the HMs-associated AS events have a significant impact on cell fate decision in a cell-cycle-dependent manner. The most intriguing discovery is that the HM-associated genes are enriched in G2/M phases and predominantly function in ATM/ATR-mediated DNA response. Evidentially, the ATM/ATR-mediated checkpoint has been recently revealed to attenuate pluripotency state dissolution and serves as a gatekeeper of the pluripotency state through the cell cycle [56]. The cell cycle has been considered the hub machinery for cell fate decision [21] since all commitments will go through the cell-cycle progression. Our study expanded such machinery by linking the HMs and AS regulation to cell-cycle pathways and TFs, which together, contribute to cell fate decision (Additional file 1: Figure S10H).

We also exemplified our hypothesized mechanism by an H3K36me3-regulated isoforms switch of PBX1. In addition to its well-known functions in lymphoblastic leukaemia [78-81] and a number of cancers [82-91], PBX1 was also found to promote hESC self-renewal by corporately binding to the regulatory elements of *NANOG* with KLF4 [101]. We found that the transcriptions of two isoforms of *PBX1, PBX1a* and *PBX1b*, are regulated by H3K36me3 during hESC differentiation. Their protein isoforms competitively bind *NANOG* and the binding of PBX1b will abolish the binding of PBX1a, which further attenuates the activity of the core pluripotency regulatory network composed of Yamanaka factors. The switch from PBX1a to PBX1b is modulated by H3K36me3 via the PSIP1/SRSF1 adapter system [47]. Our results were also supported by an extended dataset of 56 cell lines/tissues from the Roadmap/ENCODE projects. Collectively, our findings expanded understanding of the core transcriptional network by adding a regulatory layer of HM-associated AS (Figure 6F).

A very recent report showed that the switch in Pbx1 isoforms was regulated by Ptbp1 during neuronal differentiation in mice [106], indicating a contradiction that the AS of PBX1 should be negatively regulated by H3K36me3 via the MRG15/PTBP1 [46]. Our study also included the neuronal lineage and showed that differentiation to NPC is an exception, distinct from other lineages. If NPC is considered separately, the results are consistent with the recent report [106] showing that NPCs and mature neurons express increasing levels of PBX1a rather than PBX1b (Additional file 1: Figure S9B). Another recent report showed that PBX1 was a splicing target of SRSF1 in the HeLa cell line [69], which strongly supports our findings. Taken together, these evidence suggests that there are two parallel mechanisms regulating PBX1 isoforms in embryonic development, in which neuronal differentiation adopts a mechanism that is different from other lineages.

Finally, it is worth noting that both our work and other studies [68] reported that HMs cannot explain all AS events identified either during ESC differentiation or based on pairwise comparisons between cell types. Moreover, bidirectional communication between HMs and AS has been widely reported. For instance, the AS can enhance the recruitment of H3K36 methyltransferase HYPB/Set2, resulting in significant differences in H3K36me3 around the AS exons [68]. These findings increased the complexity of defining the cause and effect between HMs and AS. Nevertheless, our findings suggest that at least a subset of AS events are regulated epigenetically, similar to the way that epigenetic states around the transcription start sites define what parts of the genome are expressed. Additionally, as we described in our previous study, the AS outcomes may be estimated more precisely by combining splicing *cis-* elements and *trans*-factors (i.e., genetic splicing code) and HMs (i.e., epigenetic splicing code), as an 'extended splicing code' [20]. Taken together, we presented a mechanism conveying the HM information into cell fate decision through the AS of cell-cycle factors or the core components of pathways that controlling cell-cycle progression (Additional file 1: Figure S10H).

## Conclusions

We performed integrative analyses on transcriptome and epigenome data of the hESCs, H1 cell line, and five differentiated cell types, demonstrating that three of 16 HMs were strongly associated with half of AS events upon hESC differentiation. We proposed a potential epigenetic mechanism by which some HMs contribute to ESC fate decision through the AS regulation of specific pathways and cell-cycle genes. We experimentally validated one cell cycle-related transcription factor, PBX1, which demonstrated that alternative splicing provides a mechanism conveying the histone modification information into the regulation of cell fate decisions (Figure 6F). Our study will have a broad impact on the field of epigenetic reprogramming in mammalian development involving splicing regulations and cell cycle machinery.

## Methods

### Identification of alternative splicing exons upon hESC differentiation

Aligned BAM files (hg18) for all six cell types (H1, ME, TBL, MSC, NPC, and IMR90) were downloaded from the source provided in reference [40]. Two BAM files (replicates) of each cell type were analyzed using rMATS (version 3.0.9) [58] and MISO (version 0.4.6) [107]. The rMATS was used to identify AS exons based on the differential' per cent splice in' (PSI, ψ) values between each differentiated cell type and H1 cells (Additional file 1: Figure S1B). The splicing changes (∆PSIs or ∆ψ) are used to identify the AS events between H1 and other cell types. A higher cutoff is always helpful in reducing the false positive while compromising the sensitivity. The cutoff, |∆ψ| ≥ 0.1 or |∆ψ| ≥ 10% is widely accepted and used in AS identification [26, 108-110]. Many other studies even used 0.05 as the cutoff [111-115]. We did additional correlation analyses based on different ∆PSI cutoffs (0.1, 0.2, 0.3, 0.4, and 0.5). With the increase of the cutoffs, the number of AS events was significantly reduced (Additional file 1: Figure S11A), however, the correlations ware only slightly increased between AS and some histone modifications (Additional file 1: Figures S11B-C, upper panels), i.e. no consistently impacts of the cutoffs on the correlations were observed. Similarly, the correlation significances were also not consistently affected (Additional file 1: Figure S11B-C, lower panels). Therefore, in our study, only AS exons which hold the |∆PSI| ≥ 0.1, p-value ≤ 0.01, and FDR ≤ 5%, were considered as final hESC differentiation-related AS exons.

All identified AS event types are summarized in Additional file 1: Table S1. Finally, two types of AS exons, namely skipped exons (SE) and mutually excluded exons (MXE), which are most common and account for major AS events, were used for subsequent analysis (Additional file 2: Table S2). MISO was used to estimate the confidence interval of each PSI and generate Sashimi graphs [116] (see Figures 1A, B and 5A). To match the ChIP-seq analysis, genomic coordinates of identified AS events were lifted over to hg19 using LiftOver tool downloaded from UCSC.

### ChIP-seq data process and histone modification profiling

ChIP-seq data (aligned BAM files, hg19) were downloaded from Gene Expression Omnibus (GEO, accession ID: GSE16256) [40]. This dataset includes the ChIP-seq reads of up to 24 types of HMs for six cell types (H1, ME, TBL, MSC, NPC, and IMR90). Among these, nine histone acetylation modifications (H2AK5ac, H2BK120ac, H2BK5ac, H3K18ac, H3K23ac, H3K27ac, H3K4ac, H3K9ac, and H4K8ac) and seven histone methylation modifications (H3K27me3, H3K36me3, H3K4me1, H3K4me2, H3K4me3, H3K79me1, and H3K9me3) are available for all six cell types and therefore were used for our analyses.

To generate global differential profiles of histone modification changes between AS exons and constitutive exons upon hESC differentiation, for each MXE and SE AS events, we first randomly selected the constitutive splicing (CS) exons from the same genes, composing a set of CS exons. We then considered the histone modification changes in a ±150bp region flanking both splice sites of each AS and CS exon, i.e. a 300bp exon-intron boundary region. Each region was 15bp-binned. Alternatively, for a few cases where the exon or intron is shorter than 150 bps, the entire exonic or intronic region was evenly divided into 10 bins. This scaling allows combining all regions of different lengths to generate uniform profiles of histone modification changes around the splice sites (see Figures 2C and Additional file 1: Figure S4). To this end, we calculated the sequencing depth-normalized ∆ reads number for each binned region between H1 cells and differentiated cells as follows:

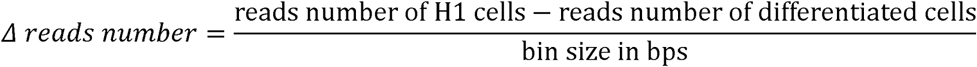

Each region is assigned a value representing the average ∆ reads number between H1 cells and differentiated cells for each histone modification. We also compared histone modification profiles between the inclusion-gain and -loss exons (Figure 3A and Additional file 1: Figure S5B) using the same strategy. The statistical results (Figure 2C and Additional file 1: Figure S5B) are presented as the p-values based on Mann-Whitney-Wilcoxon tests (using the R package).

To quantitatively link HMs to alternative splicing upon hESC differentiation, the ChIP-seq data were further processed by narrow peak calling. For each histone ChIP-seq dataset, the MACS v2.0.10 peak caller (https://github.com/taoliu/MACS/) [67, 117] was used to compare ChIP-seq signal to a corresponding whole cell extract (WCE) sequenced control to identify narrow regions of enrichment (narrow peak) that pass a Poisson p-value threshold of 0.01. All other parameters (options) were set as default. We then compared the histone modification signals between H1 cells and differentiated cells. We defined the 'differential HM signals (∆HMs)' as the difference of the normalized peak signals (i.e., the heights of the narrow peaks) between H1 and the differentiated cells. Because the 3' splice sites (3' ends of the introns) determine the inclusion of the downstream AS exons [118] and the distances from the peaks to their target sites affect their regulatory effects [119], we normalize the peak signals against the distance (in kb) between the peak summits and 3' splice sites (Additional file 1: Figure S5A). Since there is no evidence showing that distal HMs could regulate the AS, we only considered local peaks with at least one bp overlapping on either side of the AS exon. For exons without overlapping peaks, peak signals of these exons were set to zero. For the exons there are more than one overlapping peaks, the peak signals of these exons were set to the greater ones. For MXE events, only the upstream AS exons were considered due to their exclusiveness in inclusion level between these two exons, i.e., the sum of the PSIs for two exons of an MXE event is always one.

### Association studies and k-means clustering

To quantitatively estimate associations between HMs and AS, we first used three independent correlation tests, including Pearson correlation (PC), multiple linear regression (MLR), and logistic regression (LLR), to test global correlations between AS events and each of 16 HMs based on differential inclusions (∆PSIs) and differential HM signals (∆HMs). PC was performed using the R package (stats, cor.test(),method = 'pearson'). MLR and LLR were calculated using Weka 3 [120], wherein the ∆HMs are independent variables and ∆PSIs are response variables. The results show that only some HMs correlate with AS, and most correlations are weak (Additional file 1: Figure S5C). HMs that have significant correlations (p ≤ 0.05) with AS were used for further clustering analysis, through which we identified six subsets of HM-associated AS events (Additional file 3: Table S3).

*K*-means clustering was performed separately on inclusion-gain and inclusion-loss AS events of MXE and SE, based on the selected HM signatures (Additional file 1: Figure S5C, checked with a “V”). *K* was set to six for all clustering processes (Additional file 1: Figure S5D), which produced the minimal RSME (root mean square error) for most cases based on a series of clustering with *k* varying from two to eight (data not shown). Then the two clusters that generate mirror patterns, of which one from inclusion-gain events and one from inclusion-loss events, were combined to be considered as a subset of HM-associated AS events (Additional file 1: Figure S6). Finally, we identified six subsets of HM-associated AS events displaying significantly positive or negative correlations with three HMs, respectively.

### Gene expression quantification

For each cell type, two aligned BAM files (replicates) were used to estimate the expression level of genes using Cufflinks [121]. Default parameters (options) were used. The expression level of each gene was presented as FPKM for each cell type. Differentially expressed genes (DEGs) were defined as those genes whose fold changes between the differentiated cells and hESCs are greater than 2. Specifically for DEG analysis of SF genes, we collected a list of 47 ubiquitously expressed SFs with "RNA splicing" in their GO annotations from AmiGO 2 [122]. The enrichment significances in Additional file 1: Figures S1D and S3H are shown as the p-values based on hypergeometric tests, using the DEGs of all expressed genes (with the FPKM ≥ 1 in at least hESCs or one of the differentiated cell types) as background. We found that the AS genes are generally not differentially expressed between hESCs and differentiated cells, indicating that they function in hESC differentiation via isoform switches rather than expression changes. Few SF genes show differential expression between hESCs and differentiated cells, indicating the existence of epigenetic control of AS, rather than the direct control on SFs expression.

### Genome annotations

Since the RNA-seq reads (BAM) files and ChIP-seq read (BAM) files downloaded from the public sources were mapped to different human genome assemblies, NCBI36/hg18 (Mar. 2006) and GRCh37/hg19 (Feb. 2009), respectively, we downloaded two version of gene annotations (in GTF formats) from the UCSC Table Browser [123]. The hg18 GTF file was used for rMATS and MISO to identify AS during the differentiation from H1 ESCs into five differentiated cells. The hg19 GTF file was used to define the genome coordinates of AS exons and further for ChIP-seq profile analysis (Figures 2A-C, 3A, and Additional file 1: Figures S4 and S5A). We compared exonic and intronic lengths based on hg18 annotation (Additional file 1: Figure S2).

### Gene ontology enrichment analysis

The Gene Ontology (GO) enrichment analysis was performed using ToppGene [124] by searching the HGNC Symbol database under default parameters (p-value method: probability density function). Overrepresented GO terms for the GO domain 'biological process' were used to generate data shown in Figures 4C-E and Additional file 1: Figures S8A-C using either the FDR (0.05) adjusted p-value or the enriched gene numbers (Additional file 1: Figure S8A).

### Canonic pathway enrichment analysis

Both the HM-associated (n = 282) and –unassociated (150) AS genes from the top enriched GO term (GO:0007049) were used to perform canonic pathway enrichment (Figure 4F and Additional file 1: Figure S8D) analysis through Ingenuity Pathway Analysis (IPA, http://www.ingenuity.com/products/ipa).

### Stemness signature and TF binding enrichment analysis

StemChecker [61], a web-based tool to discover and explore stemness signatures in gene sets, was used to calculate enrichment of AS genes in stemness signatures. Significances were tested (hypergeometric test) using the gene set from human *(Homo sapiens)* of this database as the background. For all AS genes (n = 3,865), 2,979 genes were found in this data set. 1,395 of 2,979 genes were found in the stemness signature gene set, most of which (n = 813) are ESC signature genes. Additional file 1: Figure S3A shows the enrichment significance as the -log10 p-values (Bonferroni adjusted). For HM-associated AS genes (n = 2,125), 1,992 genes were found in this data set. 966 of 1,992 genes were found in the stemness signature gene set, most of which (n = 562) are ESC signature genes. For HM-unassociated genes (n = 1,057), 987 genes were found in this data set. 429 of 987 genes were found in the stemness signature gene set, most of which (n = 251) are ESC signature genes. The significances are shown as - log10 p-values (Bonferroni adjusted) in Figure 4A.

FunRich (v2.1.2) [125, 126], a stand-alone functional enrichment analysis tool, was used for TF binding enrichment analysis to get the enriched TFs that may regulate the query genes. The top six enriched TFs of HM-associated and –unassociated AS genes are presented and shown as the proportion of enriched AS genes. It shows that HM-associated AS genes are more likely to be regulated by TFs involved in cell differentiation and development, while the HM-unassociated AS genes are more likely to be regulated by TFs involved in cell proliferation and renewal (Figure 4B).

### Roadmap and ENCODE data analysis

All raw data are available from the GEO accession IDs GSE18927 and GSE16256. The individual sources of RNA-seq data for 56 cell lines/tissues from Roadmap/ENCODE projects were listed in Additional file 4: Table S4. The RNA-seq data (BAM files) were used to calculate the PSI of the exon 7 for PBX1 in each cell line/tissue and to estimate the expression levels of all gene (FPKM), based on aforementioned strategies. The relative expression levels of PBX1a and PBX1b shown in Figure 5 and Additional file 1: Figure S10 were calculated as the individual PFKM value of each divided by their total FPKM values.

### Statistical analyses and tests

Levels of significance were calculated with Mann-Whitney-Wilcoxon test for Figures 2C, 2D, 3A, and Additional file 1: Figures S4 and S5B, with Fisher's exact test for Figures 1D, 2D, and Additional file 1: Figure S5B, with Student’s t-test for Figures 5D and Additional file 1: Figure S7A, and with a Hypergeometric test for Additional file 1: Figures S1D and S3H. Levels of correlation significance were calculated with Pearson correlation (PC), multiple linear regression (MLR), and logistic regression (LLR) for Figures 3D, 3C, and Additional file 1: Figure S5C. MLR and LLR were performed using Weka 3 [120], whereas all other tests were performed using R packages. The p-values for the enrichment analyses (Figure 4, Additional file 1: Figures S3A and S8) were adjusted either by PDR or Bonferroni (refer to the corresponding method section for details). The statistical analyses of the ChIP, RIP and western blotting assays were shown in Additional file 1: Figures S10E-G. Three replicates were conducted for each assay and the quantitatively statistical analyses were performed on the relative band intensities normalized by control genes (ß-Actin) or input signals. ImageJ (https://imagej.nih.gov/ij/) was used to quantify the band intensities. ANOVA was used for statistical significance tests.

### Cell culture and MSCs induction

Human embryonic stem cell (H1cell line) line was purchased from the WiCell Research Institute (product ID: WA01). H1 cells were cultured on the matrigel-coated 6-well culture plates (BD Bioscience) in the defined mTeSR1 culture medium (STEMCELL Technologies). The culture medium was refreshed daily. The H1-derived mesenchymal stem cells (H1-MSC) were differentiated from H1 cells as described previously [127]. Briefly, small H1 cell aggregates were cultured on a monolayer of mouse OP9 bone-marrow stromal cell line (ATCC) for 9 days. After depleting the OP9 cell layer, the cells were then cultured in semisolid colony-forming serum-free medium supplemented with fibroblast growth factor 2 and platelet-derived growth factor BB for 2 weeks. The mesodermal colonies were selected and cultured in mesenchymal serum-free expansion medium with FGF2 to generate and expand H1-MSCs. hAT-MSCs were derived from the subcutaneous fats provided by the National Disease Research Interchange (NDRI) using the protocol described previously [128]. Briefly, the adipose tissue was mechanically minced and digested with collagenase Type II (Worthington Bio, Lakewood, NJ). The resulted single cell suspension was cultured in α-minimal essential medium with 5% human platelet lysate (Cook Regentec, Indianapolis, IN, USA), 10 μg/ml gentamicin (Life Technologies, CA, USA), and 1x Glutamax (Life Technologies). After reaching ~70% confluence, the adherent cells were harvested at the passage 1 (P1) hAT-MSC/AEY239. hBM-MSCs were isolated from commercially available fresh human bone marrow (hBM) aspirates (AllCells, Emeryville, CA) and expanded following a standard protocol [129]. Briefly, hBM-MSC were cultured in α-minimal essential medium supplemented with 17% fetal bovine serum (FBS; lot-selected for rapid growth of MSC; Atlanta Biologicals, Norcross, GA), 100 units/ml penicillin (Life Technologies, CA, USA), 100 μg/ml streptomycin (Life Technologies), and 2 mM L-glutamine (Life Technologies). After reaching ~70% confluence, the adherent cells were harvested at the passage 1 (P1) hBM-MSC-5204(G). The human IMR-90 cell line was purchased from the American Tissue Type Culture Collection (Manassas, VA) and cultured in Eagle's Minimum Essential Medium (Thermo Scientific, Logan, UT) supplemented 10% fetal calf serum, 2 mM L-glutamine, 100U/mL penicillin, and 100ug/mL streptomycin at 37°C in humidified 5% CO_2_.

### RNA isolation and reverse transcription PCR (RT-PCR)

Total RNA was extracted from cells using the RNeasy Mini plus kit (Qiagen, Valencia, CA) according to the manufacturer's instructions. RT reactions for first-strand cDNA synthesis were performed with 2μg of total RNA in 25uL reaction mixture containing 5 μl of 5X first-strand synthesis buffer, 0.5 mM dNTP, 0.5 μg oligo(dT)12-18mer (Invitrogen), 200 units of M-MLV reverse transcriptase (Promega, Madison, WI) and 25 units of RNase inhibitor (Invitrogen). The mixture was then incubated at 42°C for 50 min and 52°C for 10 min. The reaction was stopped by heating at 70°C for 15 min. The PCR amplifications were carried out in 50 μl reaction solution containing 1 μl of RT product, 5 μ1 of 10X PCR buffer, 0.15 mM MgCl_2_, 0.2 mM dNTP, 0.2 mM sense and antisense primers, and 2.5 U Taq polymerase (Bohringer, Mannheim, Germany). The sequences for the upstream and downstream primers of PBX1 and β-actin are listed in Additional file 1: Table S5. PCR reaction solution was denatured initially at 95°C for 3 min, followed by 35 cycles at 95°C for 1 min, 55°C for 40 s, 72°C 40 s. The final extension step was at 72°C for 10 min. The PCR products were resolved in a 2% ethidium bromide-containing agarose gel and visualized using ChemiDoc MP Imager (Bio-Rad).

### Quantitative Real-time PCR (qRT-PCR)

The qPCR amplification was done in a 20uL reaction mixture containing 100 ng of cDNA, 10 uL 2 X All-in-One™ qPCR mix (GeneCopoeia, Rockville, MD), 0.3 mM of upstream and downstream primers and nuclear-free water. The PCR reaction was conducted with 1 cycle at 95 °C for 10 min, 40 cycles at 95 °C for 15 s, 40 °C for 30 s and 60 °C for 1 min, followed by dissociation curve analysis distinguishing PCR products. The expression level of a gene was normalized with the endogenous control gene β-actin. The relative expressions of genes were calculated using the 2-∆∆CT method, normalized by β-actin and presented as mean ± SD (n = 3) (Figure 6A). The sequences of the paired sense and antisense primers for human SOX2, NANOG, OCT4, c-MYC, KLF4 and β-actin are listed in the Additional file 1: Table S5.

### Western blotting

Cells were lysed with 1 X RIPA buffer supplemented with protease and phosphatase inhibitor cocktail (Roche Applied Science, Indianapolis, IN) and stored in aliquots at - 20 °C until use. Twenty micrograms of cell lysates were mixed with an equal volume of Laemmli sample buffer, denatured by boiling and separated by SDS-PAGE. The separated proteins were then transferred to PVDF membranes (Bio-Rad, Hercules, CA). The membranes were blocked using 5% BSA for 1 h at room temperature and incubated with the 1^st^ antibodies against in 1% BSA overnight. SOX2, OCT4, NANOG, KLF4, c-MYC and β-actin antibodies were from Cell signalling technology (Beverly, MA). PBX1 antibody was from Abcam (Cambridge, MA) and PBX1b antibody from Santa Cruz technology (Dallas, Texas). After incubated with IgG horseradish peroxidase-conjugated secondary antibodies (Cell signalling) for 2 h at room temperature, the immunoblots were developed using SuperSignal West Pico PLUS Chemiluminescent reagent (Thermo Fisher Scientific, Waltham, MA) and imaged using ChemiDoc MP Imager (Bio-Rad).

### Co-immunoprecipitation (Co-IP)

Co-IP was used to validate the PSIP1-SRSF1 interaction (Figure 6E). Cells were lysed with ice-cold non-denaturing lysis buffer containing 20 mM Tris HCl (pH 8), 137 mM NaCl, 1% Nonidet P-40, 2 mM EDTA and proteinase inhibitor cocktails. 200 μg protein were pre-cleared with 30uL protein G magnetic beads (Cell signalling) for 30 minutes at 4 °C on a rotator to reduce non-specific protein binding. The pre-cleared lysate was immunoprecipitated with the 5μL anti-SRSF1 antibody (Invitrogen) overnight and then incubated with protein G magnetic beads for 4 hours at 4 °C. Beads without anti-SRSF1 antibody were used as an IP control. The protein G magnetic beads were washed 5 times with lysis buffer and the precipitated protein collected. The SRSF1-bound PSIP1 protein (also known as LEDGF) level was determined with the PSIP1 antibody (Human LEDGF Antibody, R&D systems) using Western blot as described above. Three specific bands (one for p75 and two for p52) were detected (Figure 6E) as indicated in R&D systems website (https://www.rndsystems.com/products/human-ledgf-antibody-762815mab3468).

### Chromatin immunoprecipitation (ChIP)

ChIP assay was performed using a SimpleChIP Enzymatic Chromatin IP Kit (Cell Signaling Technology) according to the manufacturer's instruction. Briefly, 2 × 10^7^ cells were cross-linked with 1% formaldehyde and lysed with lysis buffer. Chromatin was fragmented by partial digestion with Micrococcal Nuclease Chromatin. The protein-DNA complex was precipitated with ChIP-Grade Protein G Magnetic Beads (Cell signalling) and ChIP-validated antibodies against H3K36me3 (Abcam), PSIP1 (Novus), and PBX1b (Santa Cruz). Normal mouse IgG and normal rabbit IgG were used as negative controls. After reversal of protein-DNA cross-links, the DNA was purified using DNA purification spin columns. The purified DNA fragments were then amplified with the appropriate primers on T100 thermal cycler (Bio-Rad). The primer pairs used for PCR are listed in the Additional file 1: Table S5. The H3K36me3- and PSIP1-immunoprecipitated DNA fragments surrounding the exon 7 of PBX1b and the promoter region of NANOG in the PBX1b-immunoprecipitated DNA fragments were PCR-amplified. The ChIP-PCR products were revealed by electrophoresis on a 2% agarose gel (Figure 6C).

### RNA immunoprecipitation (RIP)

RIP assay was performed using Magna RIP™ RNA-Binding Protein Immunoprecipitation Kit (Millipore Sigma) following the manufacturer's instruction. Briefly, the cells were lysed with RIP lysis buffer with RNase and protease inhibitors. 500 μg of total protein was precleared with protein G magnetic beads for 30 minutes. The protein G magnetic beads were preincubated with 5 μg of mouse monoclonal anti-SRSP1 antibody or normal mouse IgG for 2 hours at 4° C. The antibody-coated beads were then incubated with precleared cell lysates at 4° C overnight with rotation. The RNA/protein/beads conjugates were washed 5 times with RIP was buffer and the RNA-protein complexes were eluted from the protein G magnetic beads on the magnetic separator. The SRSP1-bound RNA was extracted using acid phenol-chloroform and precipitated with ethanol. The RNA was then reverse-transcribed and the expression levels of PBX1a and PBX1b in the immunoprecipitated and non-immunoprecipitated (input) samples were analyzed using RT-PCR (Figure 6D).

**Abbreviations**
ESC: embryonic stem cell
AS: alternative (pre-mRNA) splicing
CS: constitutive splicing
HM: histone modification
TF: transcription factor
SF: splicing factor
MXE: mutually excluded exon
SE: skipped exon
PSI: percent splice in
H2AK5ac: H2A acetylated lysine 5
H2BK120ac: H2B acetylated Iysine120
H2BK5ac: H2B acetylated lysine 5
H3K18ac: H3 acetylated lysine 18
H3K23ac: H3 acetylated lysine 23
H3K27ac: H3 acetylated lysine 27
H3K4ac: H3 acetylated lysine 4
H3K9ac: H3 acetylated lysine 9
H4K8ac: H4 acetylated lysine 8
H3K27me3: H3 tri-methylated lysine 27
H3K36me3: H3 tri-methylated lysine 36
H3K4me1: H3 mono-methylated lysine 4
H3K4me2: H3 di-methylated lysine 4
H3K4me3: H3 tri-methylated lysine 4
H3K79me1: H3 mono-methylated lysine 79
H3K9me3: H3 tri-methylated lysine 9.

## Declarations

### Ethics approval and consent to participate

Not applicable.

### Consent for publication

Not applicable.

### Availability of data and materials

All RNA-seq and 16 HMs ChIP-seq data of H1 and five other differentiated cells are available in Gene Expression Omnibus (GEO) under accession number GSE16256 [40]. The BAM files of the RNA-seq data (two replicates for each, aligned to human genome hg18) are alternatively available at http://renlab.sdsc.edu/differentiation/download.html. Both RNA-seq and ChIP-seq data of 56 cell lines/tissues from the Roadmap/ENCODE projects [99, 100] are available on their official website (RoadMap: ftp://ftp.ncbi.nlm.nih.gov/pub/geo/DATA/roadmapepigenomics/by_sample/: ENCODE:ftp://hgdownload.cse.ucsc.edu/goldenPath/hg19/encodeDCC/) and all raw files are also available at GEO from the accession IDs GSE18927 and GSE16256. Additional file 4: Table S4 provides the detailed information of these data.

## Competing interests

The authors declare that they have no competing interests.

### Funding

This work was funded by the National Institutes of Health (NIH) [1R01GM123037 and AR069395].

### Author’s Contributions

YX, WZ, and XZ conceived the study. YX carried out the sequencing data analysis and interpreted the results. WZ conducted the experimental validations. WZ, SO, and KP worked on growing and characterizing the MSCs. YX wrote the first draft of the manuscript and all authors revised it. All authors read and approved the final version of the manuscript.

## Acknowledgements

We would like to give our special thanks to Drs Guangxu Jin, Liang Liu, and Dongmin Guo from Wake Forest School of Medicine, who gave a lot of comments and suggestions on the data analyses and interpretations. We also acknowledge the editorial assistance of Karen Klein, MA, in the Wake Forest Clinical and Translational Science Institute (UL1 TR001420; PI: McClain).

## Additional files

**Additional file 1: Figure S1**. Identifying hESC differentiation-related AS exons. **Figure S2**. The hESC differentiation-related AS exons possess the typical properties of AS exons. **Figure S3**. AS profiles upon hESC differentiation show lineage-specific splicing pattern. **Figure S4**. Histone modifications (HMs) change significantly around the alternatively spliced (AS) exons upon hESC differentiation. **Figure S5**. A subset of AS events are significantly associated with some HMs upon hESC differentiation. **Figure S6**. K-means clustering based on selected epigenetic features of eight HMs for MXE and SE AS exons. **Figure S7**. HM-associated AS genes are more lineage-specific. **Figure S8**. HM-unassociated AS genes are enriched in G1 cell-cycle phase and pathways for self-renewal. **Figure S9**. Isoform switch from PBX1a and PBX1b during hESC differentiation. **Figure S10**. Isoform switch of PBX1 links H3K36me3 to hESC fate decision. **Figure S11**. The effect of ∆PSI cutoffs for AS-HM correlations. **Table S1**. The number of all AS events identified during hESC differentiation. **Table S5**. The PCR primers used in this study. (PDF 2.1MB)

**Additional file 2: Table S2**. Alternative splicing events (AS exons) during the differentiation from H1 cells to differentiated cells. (XLSX 1.8MB)

**Additional file 3: Table S3**. HM-associated AS exons based on k-means clustering. (XLSX 1.1MB)

**Additional file 4: Table S4**. 56 cell lines/tissues and their corresponding RNA-seq data sources from ENCODE and Roadmap projects. (XLSX 14.5KB)

